# Is N-Hacking Ever OK? A simulation-based inquiry

**DOI:** 10.1101/2019.12.12.868489

**Authors:** Pamela Reinagel

**Affiliations:** Section of Neurobiology, Division of Biological Science, University of California, San Diego

## Abstract

After an experiment has been completed, a trend may be observed that is “not quite significant”. Sometimes in this situation, researchers collect more data in an effort to achieve statistical significance. Such “N-hacking” is condemned because it can lead to an excess of false positive results. I use simulations to demonstrate how N-hacking causes false positives. However, in a parameter regime relevant for many experiments, the increase in false positives is quite modest. Moreover, results obtained this way have higher Positive Predictive Value than non-incremented experiments of the same sample size and statistical power. In other words, adding a few more observations to shore up a nearly-significant result can *increase* the reproducibility of results, counter to some current rhetoric. Many experiments are non-confirmatory, and unplanned sample augmentation with reasonable decision rules would not cause rampant irreproducibility in that context.

## Introduction

There has been much concern in recent years concerning the lack of reproducibility of results in some scientific literatures, leading to a call for improved statistical practices [1-5]. The call for better education in statistics and greater transparency in reporting is justified and welcome. But we shouldn’t apply rules and procedures by rote without comprehension. Experiments often require substantial financial resources, scientific talent, and use of animal subjects. There is an ethical imperative to use these resources efficiently. To ensure both reproducibility and efficiency of research, experimentalists need to understand statistical principles rather than blindly apply rules.

As an example of a mis-applied rule, I will explore a rule of null hypothesis significance testing: if a sample size N is chosen in advance, it may not be changed (augmented) after seeing the results [1, 6-9]. In my experience this rule is not well known among biologists, and is commonly violated. Many researchers engage in “N-hacking” – incrementally adding more observations to an experiment when a preliminary result is “almost significant”. It not uncommon for reviewers of manuscripts to require the authors collect more data to support a claim if the presented data do not reach significance. Prohibitions against collecting additional data therefore meet with considerable resistance and confusion in this community.

For the sake of accessibility to biologist readers, this paper will use only empirical demonstrations. Statistician readers may rest assured that there is nothing theoretically new here, nor am I claiming or attempting to overturn any established statistical principle. But even for those familiar with the theoretical principles at play, the numerical results may be surprising. The specific sampling heuristic simulated here is meant to be descriptive of practice, and is different in details from established formal adaptive sampling methods [6, 10-12].

The simulations in this paper can be taken to represent a large number of independent studies, each collecting separate samples to test a different hypothesis. All simulations were performed in MATLAB 2018a. Definitions of all terms and symbols are summarized in Appendix 1. The MATLAB code for all these simulations and more can be found in [13], along with the complete numeric results of all computationally intensive simulations.

### Part I: Effects of N-hacking on the false positive rate

Experiments were simulated by comparing two independent samples of size *N* drawn from the same normal distribution. An independent sample Student’s *t*-test was used to accept or reject the null hypothesis that the samples came from distributions with the same mean, with the significance threshold *p* < 0.05. Because the samples always came from the same distribution, any positive result is a false positive. I will call the observed false positive rate when the null hypothesis is true *FP*_0_ (“FP null”), also known as the Type I Error Rate, to emphasize that this is not the same as “the probability a positive result is false” (False Positive Risk). By construction the *t*-test produces false positives at a rate of exactly *α*, the significance threshold (0.05 in this case).

If researchers continue collecting more data until they get a significant effect, however, some true negatives will be converted to false positives. For example, suppose many separate labs each ran a study with sample size N=8, where in every case there was no true effect to be found. If all used a criterion of *α* = 0.05, we expect 5% to obtain false positive results. But suppose all the labs with “nonsignificant” outcomes responded by adding four more data points to their sample and testing again, repeating this as necessary until either the result was significant, or the sample size reached N=1000. The interim “*p* values” would fluctuate randomly as the sample sizes were grown (Figure 1A). In some cases, the “*p* value” would cross the significance threshold by chance. If these studies ended as soon as *p* < *α* and reported significant effects, these would represent excess false positives, above and beyond the 5% they intended to accept.

**Figure 1.**
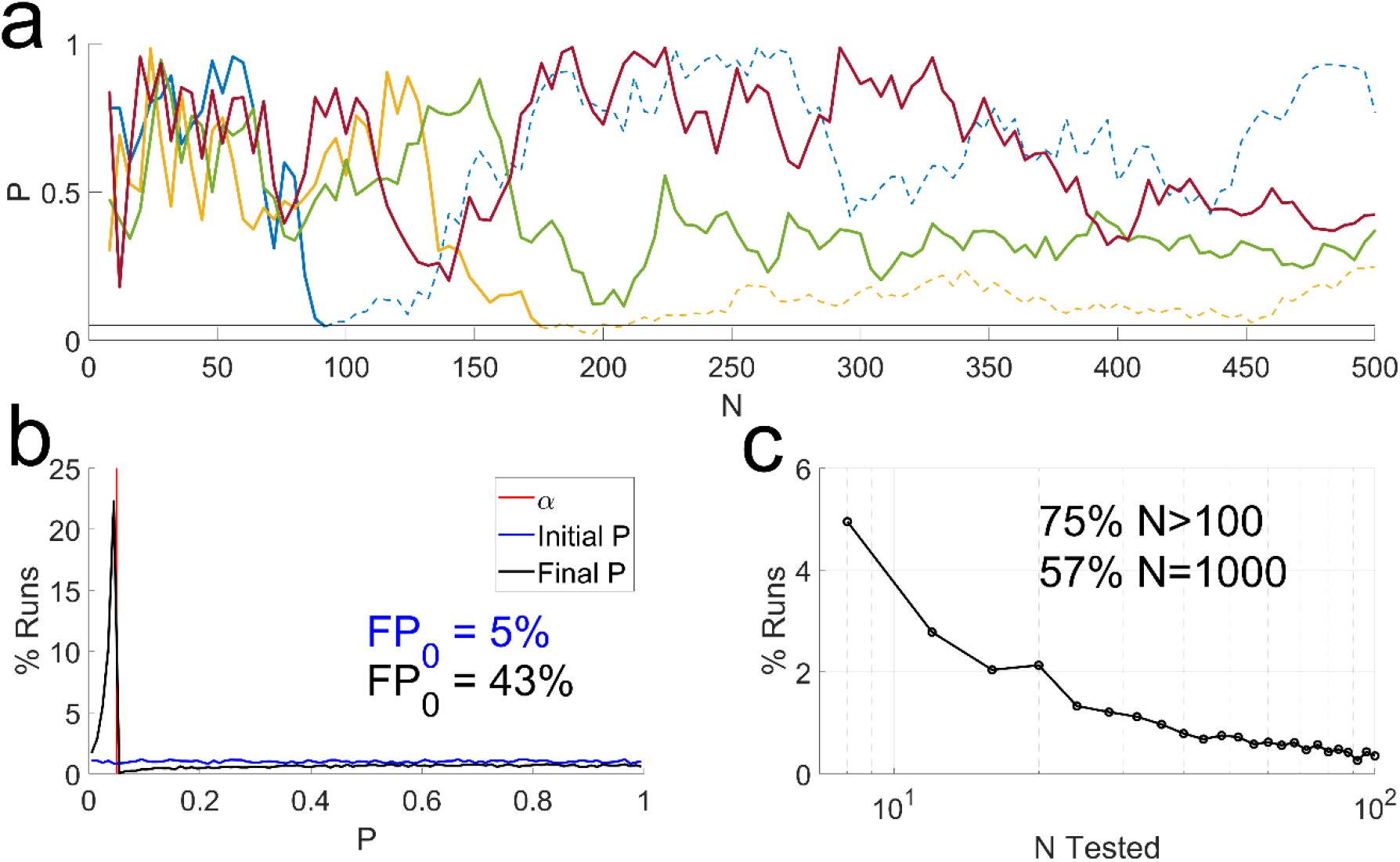
The problem with N-hacking. Simulation of experiments in which there was no true effect, starting with samples of size N=8. If the result was nonsignificant, we added 4 more and retested, until either the result was significant or N=1000. **a**. Evolution of “p values” of four simulated experiments, as N was increased. If sampling were terminated when *p* < *α* (solid blue and gold curves) this would produce false positives. If sampling had continued, those would have become nonsignificant again (dashed blue and cold curves). **b**. Distribution of initial and final “p values” of 10^5^ such experiments, in bins of width 0.01. Vertical red line indicates the nominal *α* (0.05). FP_0_ values indicate the false positive rates associated with the same colored curves (integral from *p* = 0 to *p* = α).**c**. Distribution of final sample sizes, based on counts of each discrete sample size. The fraction of runs that exceeded N=100 or that reached N=1000 are indicated.

In a simulation of 10,000 such experiments, there were 495 false positives (5%) in the initial t-test, but 4262 false positives (43%) after N-hacking (Figure 1B). Therefore, the final “*p* values” after N-hacking are not valid *p* values – they do not reflect the probability of observing a difference at least this large by chance if there were no real effect. This has been pointed out by many others [1, 6-9], and serves to illustrate why N-hacking can be problematic for users of *p* values.

This scenario postulates unrealistically industrious and stubborn researchers, however. Suppose the experimental units were mice. For the 5% of labs that obtained a false positive at the outset, the sample size was a reasonable N=8 mice. All other labs had larger final samples. Three quarters of the simulated labs would have tested over 100 mice, and over half of the simulated labs tested 1000 mice before giving up (Fig. 1C). Moreover, in 75% of the simulated runs, additional data were collected after observing an interim “*p* value” in excess of 0.9. These choices are frankly implausible.

Suppose instead that the sample size would be increased only if *p* < 0.10, and only up to a maximum of N=32 mice. In this constrained N-increasing procedure, *p* values falling within the eligible window (0.05 ≤ *p* < 0.10) are treated as inconclusive outcomes, or can be viewed as defining a “promising zone” of the negative results most likely to be false negatives. These inconclusive/promising cases are resolved by collecting additional data. Experiments with interim *p* values falling above the upper limit are considered futile and abandoned. The strict upper limit on the sample size reflects the fact that real experiments have finite resources; however, I will mostly explore procedures that rarely reach this limit.

This constrained version of N-hacking (more neutrally, “sample augmentation”) also yielded an increase in the rate of false positives, but this effect was rather modest, yielding a false positive rate *FP*_0_ = 0.0625 instead of the intended 0.05 (Figure 2a). Note that no correction for multiple comparisons was applied. Following this procedure resulted in a negligible increase in the sample size on average, and rarely more than twice the initially planned sample (Figure 2b).

**Figure 2.**
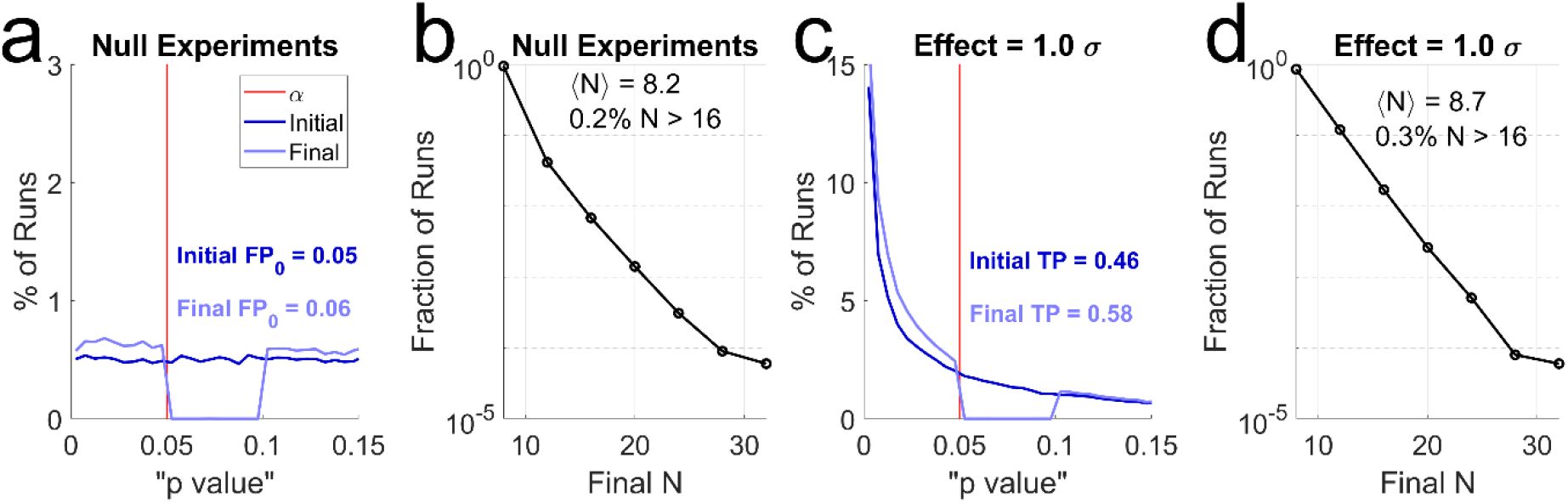
Constrained sample augmentation. Hypothetical sampling procedure in which an initial sample of *N* = 8 is incremented by 4, only if 0.05 < *p* < 0.10, up to a maximum of *N* = 32. **a**. Distribution of initial *p* values (dark blue) vs. final “*p* values” (pale blue) in simulations with no real effect. Horizontal scale is expanded in the region around *α* (red line) to show detail. Note the depletion of “p values” in the eligibility window (trough in pale curve). “FP_0_” indicates the false positive rate before (Initial) vs. after (Final) augmentation. In this simulation of 10^5^ runs, the observed false positive rate of this procedure was *FP*_0_ = 0.0625. **b**. Distribution of final sample sizes in the simulations shown in a. ⟨*N*⟩ indicates the mean final sample size; the percentage of runs exceeding N=16 is also shown. Note that the sample cap of N=32 was rarely reached. **c**. Distribution of initial and final “*p* values” for the same sampling policies as a-b, when all experiments had a real effect of size 1 SD. “TP” indicates the observed true positive rate before and after augmentation. **d**. Distribution of final sample sizes of experiments in c. Mean sample size and percent exceeding N=16 are also shown.

Therefore, if a researcher routinely collected additional data to shore up almost-significant effects, constrained by a conservative cutoff for being “almost” significant, the inflation of false positives would be inconsequential. These two extreme examples (Figure 1 vs. 2) show that from the point of view of the false positive rate *FP*_0_, sample augmentation can either be disastrous or benign depending on the details of the decision rule. I will return to the question of what parameters are compatible with reasonably limited false positive rates in Part III.

In addition to false positives, however, we must also consider the effect on false negatives. To this end, I repeated the simulations above assuming a true effect of 1 SD difference in the means. In these simulations, every positive is a true positive, and every negative is a false negative. In the initial samples of *N* = 8, the real effect was detected in some but not all experiments (Fig 2c, dark blue curve). The initial true positive rate, 46%, is simply the statistical power for a fixed sample size of *N* = 8 per group to detect a 1 SD effect.

Constrained sample augmentation increased the statistical power (Fig 2c, pale blue curve), while only slightly increasing the average final sample (Fig 2d). A fixed-N experiment using the augmentation procedure’s type I error rate for *α* (*α* = 0.0625) and N=9 had less power than the augmented procedure (56%, vs 58%). Thus, constrained sample augmentation can increase the chance of discovering real effects, even compared to fixed-N experiments with the same false positive rate and an equal or larger final sample size.

### Part II. Effects on Positive Predictive Value

What most biologists really care about, however, is whether their positive results will be reliable in the sense of identifying real effects. This is not given by 1 − *p* or 1 − *α*, as many erroneously believe, but another quantity called the positive predictive value (PPV). To determine PPV, one must also know the effect size and the fraction of experiments in which a real effect exists – the *prior probability*, or prior for short. In any real-world experiment, the true effect size and prior are unknown. But in simulations we stipulate these values, so the PPV is well-defined and can be numerically estimated.

To illustrate how sample augmentation impacts PPV, simulations were done exactly as described above, but now 10% of all experiments had a real effect (1σ, as in Fig 2c-d), and the remaining 90% had no real effect (as in Fig 2a-b). In this toy world, the point of doing an experiment would be to find out if a particular case belongs to the null group or the real effect group. The positive predictive value (PPV) is defined as the fraction of all positive results that are true positives.

#### Side Box

Positive Predictive Value.

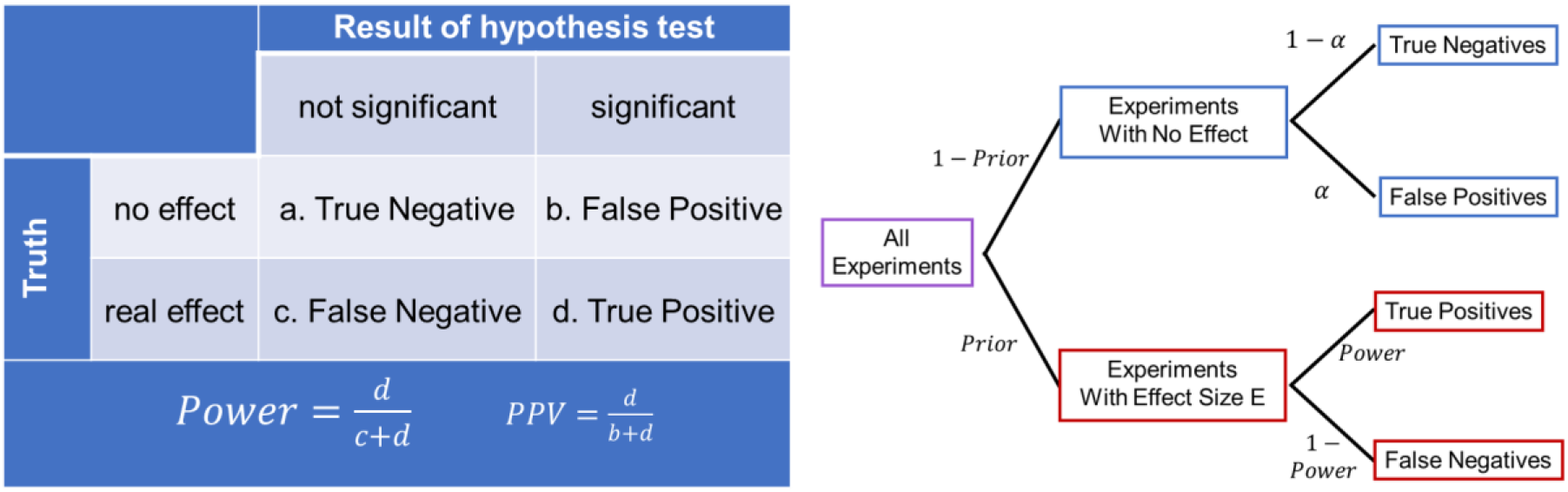

A scientific community tests many hypotheses. The effect for which any experiment is testing is in reality either absent or present (Truth, rows in table). The outcome of a binary significance test is either negative or positive (Result, columns of table). This yields four types of outcomes: a. true negatives; b. false positives (Type I errors); c. false negatives (Type II errors), and d. true positives. The statistical *power* of a procedure is defined as the fraction of real effects that yield significant effects. The *positive predictive value* (PPV) of a procedure is defined as the fraction of significant effects that are real effects. The tree diagram illustrates how these quantities are related. The probability of a false positive when there is no real effect depends only on the procedure *α* (blue boxes, upper right). The probability of a true positive when there is a real effect depends on the power (red boxes, lower right), which in turn depends on both *α* and the effect size E. The probability that a significant event is real (PPV) further depends on the fraction of all experiments that are on the red vs. blue branch of this tree, the Prior. In the real world, effect sizes and priors are not known. For a more in-depth primer see for example [14]

Constrained sample augmentation can increase both power and positive predictive value. For example, using *N* = 8 and *α* = 0.01, statistical power increased from 21% before to 28% after augmentation; PPV increased from 70% to 73%. These effects depend quantitatively on *α*, which can be shown by simulating the sampling procedure of Figure 2 for several choices of *α* (Figure 3). The average final sample size ⟨*N*_final_⟩ ranged from 8.02 (for *α* = 0.001) to 8.28 (for *α* = 0.05). Therefore, performance of constrained augmentation can be reasonably compared to the fixed-N procedure with *N* = 8 (Figure 3, red curves). The sample augmenting procedure had higher power than fixed-N, even after correcting for the false positive rate of the procedure (Figure 3a, solid blue curve), and yielded higher PPV than fixed-N, whether or not we corrected for the false positive rate of the procedure (Figure 3b).

**Figure 3.**
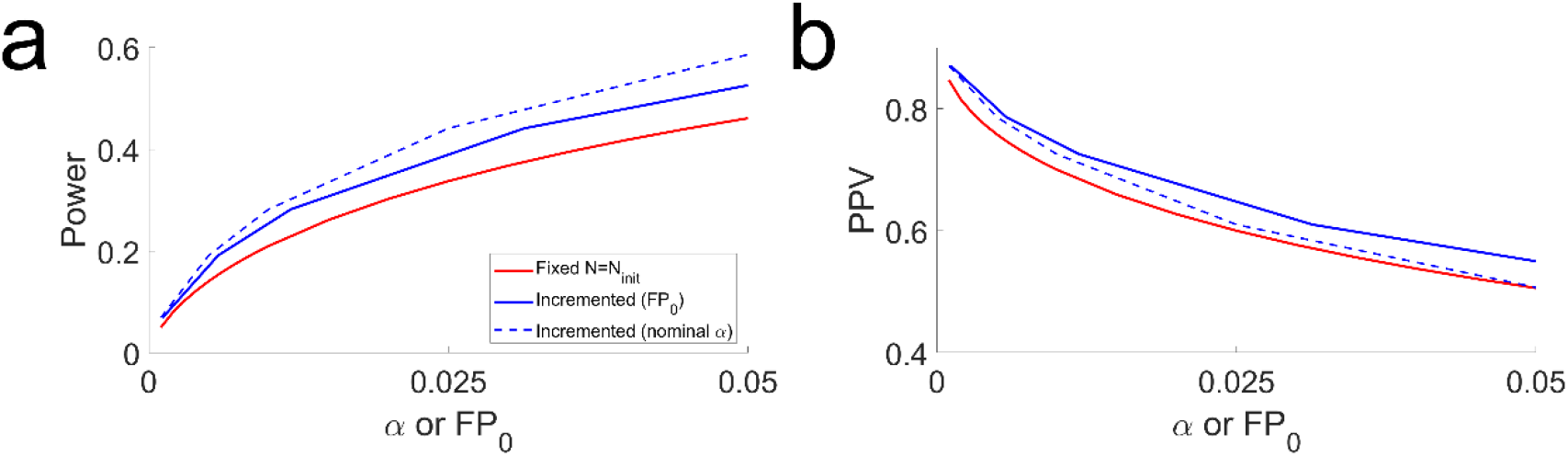
Constrained augmentation can increase both power and PPV. Simulations in which 10% of all experiments had a real effect (*Pr* = 0.1) of size 1 SD (*E* = 1σ), varying the significance criterion *α*. **a**. Statistical power of the fixed-N procedure with N=8 (red), compared to constrained augmentation with *N*_*init*_ = 8, *N*_*incr*_ = 4, *N*_*max*_ = 32, *w* = 1 (blue). For the sample augmenting procedure, results are plotted as a function of the observed false positive rate *FP*_0_ (solid blue) or the nominal criterion *α* (dashed blue). **b**. Positive Predictive Value for the same simulations analyzed in a. Statistical power and PPV were computed analytically for the fixed-N procedure, or estimated from *M* = 10^4^/*α* simulated experiments for the incrementing procedure.

In conclusion, although unplanned sample augmentation or N-hacking is widely considered a “questionable research practice”, under these conditions it would yield results at least as reliable as those obtained by sticking with the originally planned sample size, if not more reliable.

### Part III. Parameter Dependence

#### Dependence of FP_0_ on parameters

Given that unconstrained sample augmentation can increase false positives drastically (Figure 1) whereas constrained sample augmentation increases false positives negligibly (Figure 2), it would be useful to have a general rule for what false positive rate to expect for any arbitrary constraint condition. This would be quite difficult to derive analytically, but can easily be explored using numerical simulations.

The critical factor for the false positive rate *FP*_0_ is the width of the window of *p* values that are eligible for augmentation, relative to the significance criterion *α*. To express this, I define the variable *w* as the width of the eligibility window in units of *α*. For example, in the case of Fig 2a, *α* = 0.05 and *w* = 1, such that one would accept a hypothesis if *p* <0.05, reject if *p* ≥ 0.10, and add observations for the inconclusive *p* values in between. In the egregious N-hacking case simulated in Figure 1, *α* = 0.05 and *w* = 19, such that one would accept a hypothesis if *p* <0.05, reject if *p* > 1.00, and increment otherwise. For a table of the lower and upper boundary *p* values defining the inconclusive/promising window for different choices of *w*, see Supplementary Table 1. Below I will call the initial sample size of an experiment *N*_*init*_, the number of observations added between re-tests the sample increment *N*_*incr*_, and the maximum sample size one would test *N*_*max*_. A table of these and other variable definitions is provided in Appendix 1.

Before discussing simulation results, we can develop an intuition. The false positive rate after sample augmentation cannot be less than *α*, because this many false positives are obtained when the initial sample is tested. Subsequent sample augmentation can only add to the false positives. Furthermore, *FP*_0_ cannot exceed the upper cutoff *p* value of *α*(1 + *w*), because all experiments with initial *p* values above this are immediately abandoned as futile. Exactly *wα* experiments are deemed inconclusive or promising (eligible for additional data collection), so no more than this many can be converted to false positives. Indeed, no more than half of them should be converted, because if there is no true effect, collecting additional data is more likely to shift the observed effect towards the null than away from it.

To show this numerically, I simulated a range of choices of both, using *N*_*init*_ = 12, *N*_*incr*_ = 6, *N*_*max*_ = 24 (Figure 4). I focused on what I consider realistic choices of *α* (not exceeding 0.1) and *w* (not exceeding 1). In this range, simulations show that for any given choice of *w*, the false positive rate depends linearly on *α* (Figure 4a). The slopes of these lines are in turn an increasing function of the decision window *w* (Figure 4b, symbols). For small *w*, this relationship is approximately linear.

**Figure 4.**
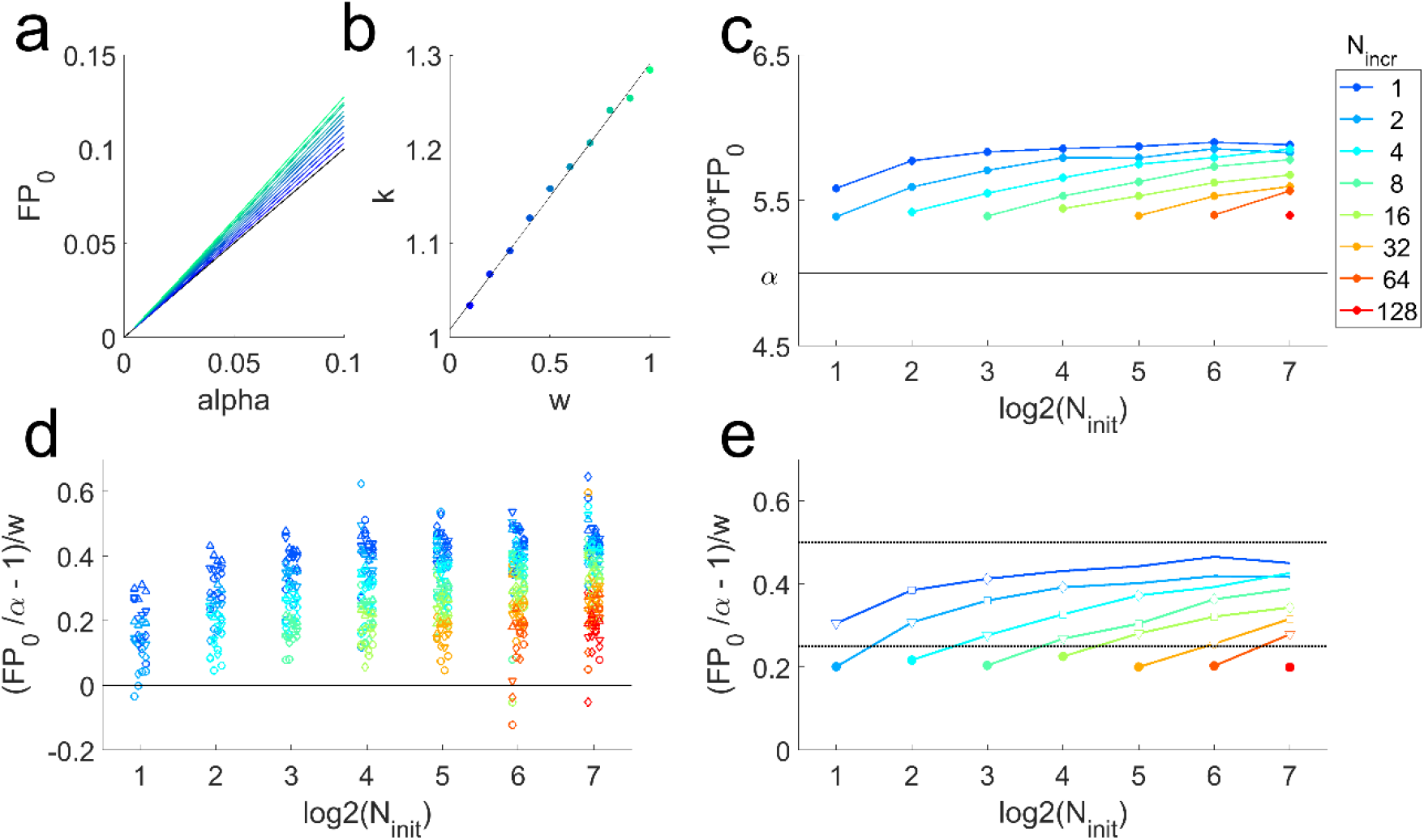
Dependence of false positive rate *FP*_0_ on α and *w*. **(a)** Simulations of constrained sample augmentation when the null hypothesis is true, using *N*_*init*_ = 12, *N*_*incr*_ = 6, *N*_*max*_ = 24, *M* = 10^6^ simulated experiments per condition. The observed false positive rate *FP*_0_ vs. *α*. Color indicates *w* (cf. panel b). For each *w, FP*_0_is plotted for each simulated value of *α* [0.005, 0.01, 0.025, 0.0500, 0.1], and the data points connected. The identity line (black), *FP*_0_ = *α*, is the false positive rate of the standard Fixed-N procedure. **(b)** The slopes *k* obtained from linear fits to the data shown in (a), plotted as a function of window size *w* (colored symbols). The dependence of the slope *k* on *w* is not linear in general, but is approximately linear in this parameter range (linear fit, black). **(c)** The realized false positive rate *FP*_0_ of the constrained N-increasing procedure, as a function of log_2_ *N*_*init*_ (horizontal axis) and *N*_*incr*_(colors), for the case *α* = 0.05, *w* = 0.4, *N*_*max*_ = 256. The false positive rate is always elevated compared to α (black line), but this is more severe when the intial sample size is larger (curves slope upward) or the incremental sample growth is smaller (cooler colors are higher). Color key at right applies to panels c-e. **(d)** Results for four choices of *α* (0.005, 0.010, 0.025, or 0.050; symbol shapes) and w (0.1, 0.2, 0.3 or 0.4, small horizontal shifts), plotted as 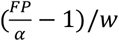 (vertical axis) to reveal regularities. For the fixed-N procedure *FP*_0_ = *α*, so this equation reduces to 0 (black line). Positive values on this scale indicate an increase in the false positive rate compared to the fixed-N procedure. **(e)**. Summary of simulations in (d) obtained by fitting the equation *FP* = (*cw* + 1)*α*, as in panel b. Symbols indicate simulations in which *N*_*incr*_ = *N*_*init*_ (closed circles), *N*_*incr*_ = *N*_*init*_/2 (open triangles), *N*_*incr*_ = *N*_*init*_/4 (open squares) and *N*_*incr*_ = *N*_*init*_/8 (open diamonds). Upper dashed black line is a proposed empirical bound 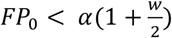. Lower black line is a proposed bound for 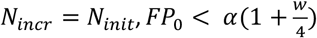.

The false positive rate also depends on the initial sample size and the increment size. To illustrate this, I repeated these simulations for *N*_*init*_ ranging from 2 to 128 initial sample points and increments *N*_*incr*_ ranging from 1 to *N*_*init*_, capping the maximum total sample size at *N*_*max*_ = 256. The false positive rate *FP*_0_ was inflated more severely when the intial sample size was larger, or the incremental sample growth step smaller (Fig 4c). The increment cannot get any smaller than *N*_*incr*_ = 1, and this curve has leveled off by *N*_*init*_ = 256, so we can take *N*_*init*_ = 256, *N*_*incr*_ = 1 (observed *FP*_0_ = 0.059) as the worst case scenario for this choice of *α* and *w*. In the explored regime, the maximum sample size was rarely reached, and therefore had little influence on overall performance characteristics.

The false positive rate is a systematic function of *α* and *w*. Because *FP*_0_ scales linearly with *α* (Fig 4a) and approximately linear with *w* over this range of *w* (Fig 4b), results of the simulations for all combinations of *α* and *w* can be summarized on one plot by linearly scaling them (Fig 4d). This confirms that the false positive rate is bounded (upper dashed line, Fig 4e), as expected from the intuition given above. When the increment step is the same as the initial sample size, there appears to be a lower bound (lower dashed line, Fig 4e). Simulations up to *w* = 0.4 are shown, but these empirically justified bounds are not violated when *w* is larger, because the dependence on *w* is sublinear, such that the *normalized* false positive rate decreases slightly as *w* increases. Of course the *absolute* false positive rate still increases with the window size *w*. In the egregious N-hacking case of Fig 1 (*α* = 0.05, *w* = 19), for example, the empirical bound yields a not-very-comforting bound of *FP*_0_ < 0.52 (still a conservative estimate relative to the numerically estimated value, *FP*_0_ = 0.42).

#### Dependence of PPV on parameters

Above I demonstrated one condition in which uncorrected sample augmentation improved both statistical power and PPV, but this is not always the case. To show this I repeated simulations like those of Figure 3 out to extreme choices of *w* (0.2 to 10), using a worst-case increment size (*N*_*incr*_ = 1) and a liberal sampling cap (*N*_*max*_ = 50), again varying *α*. The initial sample size was varied from extremely underpowered (*N*_*init*_ = 2) to appropriately powered (*N*_*init*_ = 16) for the fixed-N procedure.

For a fixed choice of *α*, increasing *w* (more aggressive sample augmentation) always increases statistical power (Figure 5 bottom row, warm colors are above cool colors along any gray curve). This makes sense: the more freely one would collect a few more data points, the more often false negatives will be rescued to true positives. However, this only sometimes increases PPV compared to the fixed-N procedure. For example, for *N*_*init*_ = 4, *α* = 0.01, PPV increases with *w* (bottom left, circles: gray curve slope is positive) but for *N*_*init*_ = 8, *α* = 0.05, PPV decreases with increasing *w* (bottom right, squares: gray curve slope is negative).

**Figure 5.**
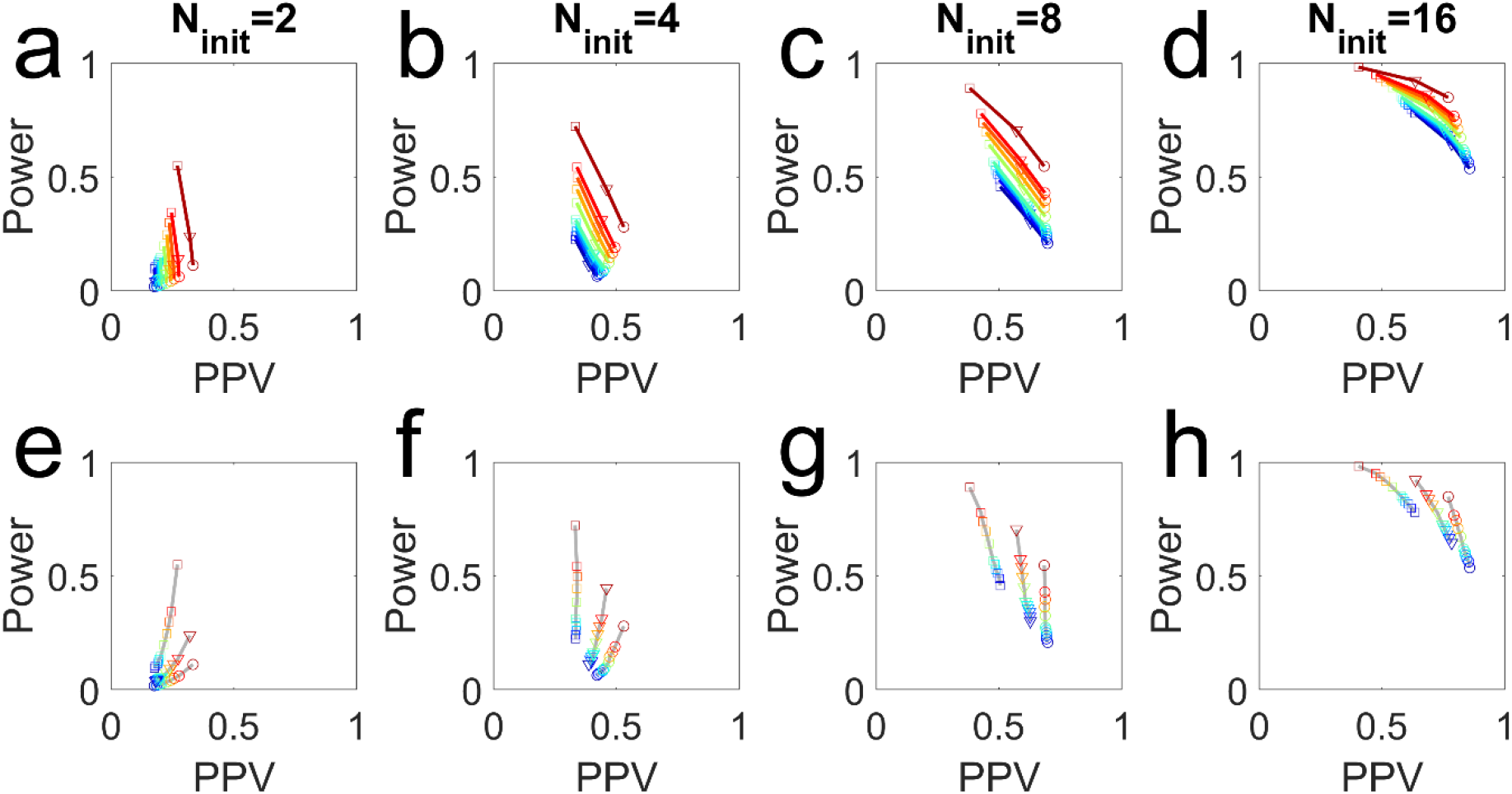
Uncorrected sample augmentation improves the FP_0_-Power or PPV-Power trade-off. Plots show the measured PPV vs. statistical power in simulations with effect size *E* = 1σ and prior effect probability *P*(*H*_1_) = 0.10, with *N*_*init*_ as indicated on column title, *N*_*incr*_ = 1, *N*_*max*_ = 50. Each symbol represents the results from *M* = 10^6^ simulated experiments, with no corrections. Symbols indicate *α* (○=0.01, ▽=0.02, □=0.05). Colors indicate *w* (blueèred= 0, 0.2, 0.4, 0.6, 0.8, 1, 2, 3, 4, 5, 10). Note that dark blue (*w* = 0) is the fixed-N procedure. **Top panels:** simulations with the same *w* and different *α* are connected with curves. **Bottom panels:** the same data, but simulations with same *α* and different *w* are connected with gray curves.

Nevertheless, uncorrected sample augmentation always produces a higher PPV and power than the fixed-N procedure with the same false positive rate; or a higher PPV and lower false positive rate than the fixed-N procedure with the same statistical power (e.g., Supplementary Table 2). For any sample-augmentation parameters (curves other than dark blue in top panels), if we find the point along the dark blue curve (Fixed-N) that has the same power, the PPV is lower; or if we find the point on the Fixed-N curve with the same PPV, the power is lower. Curves with higher *w* lie strictly above and to the right those of lower *w*, including fixed-N (*w* = 0). In this sense, N-hacking is always better than not N-hacking.

## Conclusions

Many experimental biologists are unaware that it matters when or how they decide how much data to collect when testing for an effect. The first take away of this paper is that if you are reporting *p* values, it does matter. Increasing the sample size after obtaining a non-significant *p* value will on average lead to a higher rate of false positives, if the null hypothesis is true. This has been said many times before, but most writers warn that this practice will lead to extremely high false positive rates [6-9]. This certainly *can* occur, if an experimentalist would increment their sample size no matter how far from *α* the *p* value was, and would continue collecting data until N was quite large (Figure 1). But I personally have never met an experimental biologist who would do that.

If the *p* value would have to be quite close to *α* for one to continue collecting data, the effects on the false positive rate are modest and bounded. The magnitude of the increase in the false positive rate depends quantitatively on the initial sample size *N*_*init*_, the significance criterion *α*, the promising zone or eligibility window *w*, and the increment size *N*_*incr*_. An intuitive explanation and empirical validation have been provided for an upper bound on the false positive rate. Moreover, sample augmentation strictly *increases* the positive predictive value (PPV) achievable for any given statistical power, compared to studies that strictly adhere to the initially planned N. These results were demonstrated in both underpowered and well-powered regimes. To my knowledge this particular sampling procedure has not been considered before, but the basic principles underlying the benefits of adaptive sampling have been long known in the field of statistics, for example [15].

The reproducibility crisis literature has frequently blamed optional stopping or N-hacking as an important cause of irreproducible results. But in some regimes uncorrected data-dependent sample augmentation would increase both statistical power and PPV relative to fixed-N procedure of the same nominal *α*. In research fields that operate in that restricted regime, therefore, it is simply not true that N-hacking would lead to an elevated risk of unreproducible results, as often claimed. A verdict of “statistical significance” reached in this manner is if anything *more* likely to be reproducible than results reached by fixed-N experiments with the same sample size – even if no correction is applied for sequential sampling or multiple comparisons. Therefore, if any research field operating in that parameter regime suffers from a high rate of false claims, other factors are more likely to be responsible.

### Some caveats

I have asserted that certain practices are common based on my experience, but I have not done an empirical study to support this claim. Moreover, I have simulated only one “questionable” practice: post-hoc sample augmentation based on an interim *p* value. I have seen this done to rescue a non-significant result, as simulated here; but I’ve also seen it done to verify a barely-significant one (a practice which results in *FP*_0_ < *α*). In other contexts, I suspect researchers flexibly decide when to stop collecting data based on directly observed results or visual inspection of plots, without interim statistical tests. Such decisions may take into account additional factors like the absolute effect size, a heuristic which could have even more favorable performance characteristics [16]. From a metascience perspective, a comprehensive study of how researchers make sampling decisions in different disciplines (biological or otherwise), coupled with an analysis of how the observed operating heuristics would impact reproducibility, would be quite interesting [17].

In this paper I have discussed the effect of N-hacking on Type I errors (false positives) and Type II errors (false negatives). Statistical procedures may also be evaluated for errors in effect size estimation: Type M (magnitude) and Type S (sign) errors. [18]. Even in a fixed-N experiment, effect sizes estimated from “significant” results are systematically overestimated. This bias can be quite large when N is small. This concern also applies to the low-N experiments described here, but sample augmentation did not increase either the Type M or Type S error compared to fixed-N experiments [19].

It is often necessary to collect data in batches and/or over long time periods for pragmatic reasons. Differences between batches or over time can be substantial sources of variability – even when using a fixed-N procedure. Therefore, one should check if there are batch or time-varying effects and account for them in the analysis if necessary. This is not unique to N-hacking, but with incremental sample augmentation this concern will always be applicable. Likewise, if the experimental design is hierarchical, a hierarchical model is needed, regardless of sampling procedure [20].

I have simulated a balanced experimental design, with the same N in both groups in the initial batch, and augmenting the sample size of both groups equally in each sample augmentation step. This is recommended, especially in multi-factorial designs with many groups. First, it minimizes the risk of confounding batch effects with the effects under study. Moreover, selectively augmenting the sample size in some groups but not others can introduce other confounds and interpretation complexities [21].

### So, is N-hacking ever OK?

Researchers today are being told that if they have obtained a non-significant finding with a *p* value just above *α*, it would be a “questionable research practice” or even a breach of scientific ethics to add more observations to their data set to improve statistical power. Nor may they describe the result as “almost” or “bordering on” significant. They must either run a completely independent larger-N replication, or fail to reject the null hypothesis. Unfortunately, in the current publishing climate this generally means relegation to the file drawer. Depending on the context, there may be better options.

I will use the term “confirmatory” to mean a study designed for a null hypothesis significance test, intended to detect effects supported by p values or “statistical significance”. I will use the term “non-confirmatory” as an umbrella term to refer to all other kinds of empirical research (sometimes called “exploratory” [22]). An ideal confirmatory study would completely pre-specify the sample size or sampling plan, and every other aspect of the study design; and furthermore, establish that all null model assumptions are exactly true and all potential confounds are avoided or accounted for. This ideal is unattainable in practice. Therefore, real confirmatory studies fall along a continuum from very closely approaching this ideal, to looser approximations.

A very high bar is appropriate when a confirmatory experiment is intended to be the sole or primary basis of a high-stakes decision, such as a clinical trial to determine if a drug should be approved. At this end of the continuum, the confirmatory study should be as close to the ideal as humanly possible, and public pre-registration is reasonably required. The “*p* value” obtained after unplanned incremental sampling is not a valid *p* value, because without a pre-specified sampling plan, you can never truly know or prove what you would have done if the data had been otherwise, so there is no way to know how often a false positive would have been found by chance. N-hacking forfeits control of the Type I Error rate, whether the false positive rate is increased or decreased thereby. Therefore, in a strictly confirmatory study, N-hacking is not OK.

That being said, *planned* incremental sampling is not N-hacking. There are many established adaptive sampling procedures which allow flexibility in how much data to collect while still producing rigorous *p* values. These methods are widely used in clinical trials, where costs as well as stakes are very high. It is beyond the present scope to review these methods but see [6, 10-12]. Simpler or more convenient pre-specified adaptive sampling schemes are also valid, even if they are not optimal [8]. In this spirit, the sampling heuristic I simulated could be followed as a formal procedure (Appendix 2).

A less-perfect confirmatory study is often sufficient in lower-stakes conditions, such as when results are intended only to inform decisions about subsequent experiments, and where claims are understood as contributing to a larger body of evidence for a conclusion. In this research context, transparent N-hacking in a mostly pre-specified study might be OK. Although data-dependent sample augmentation will prevent determination of an exact *p* value, the authors may still be able to estimate or bound the p value (see Appendix 2). When such a correction is small and well justified, this imperfection might be on a par with others we routinely accept, such as assumptions of the statistical test which cannot be confirmed or which are only approximately true.

The reader is cautioned that the following opinion is controversial, but <u>with full disclosure</u>, I think it is acceptable to report a *p* value in this situation. The report should disclose that unplanned sample augmentation occurred, report the interim *N* and *p* values, describe the basis of the decision as honestly as possible, and provide and justify the authors’ best or most conservative estimate of the *p* value. With complete transparency (including publication of the raw data), consumers of the study can decide what interpretation of the data is most appropriate for their purposes - including relying only on the initial, strictly confirmatory *p* value, if that standard is most appropriate for the decision they need to make.

It must be mentioned, however, that many high-quality research studies are mostly or entirely non-confirmatory, even if they follow a tightly focused trajectory or are hypothesis (theory) driven. For example, “exploratory experimentation” aims to describe empirical regularities prior to formulation of any theory [23]. Development of a mechanistic or causal model may proceed through a large number of small (low-power) experiments [24, 25], often entailing many “micro-replications” [26]. In this type of research putative effects are routinely re-tested in follow-up experiments or confirmed by independent means [27-30]. Flexibility may be essential to efficient discovery in such research, but the interim decisions about data collection or other aspects of experimental design may be too numerous, qualitative, or implicit to model. In this kind of research, the use of *p* values is entirely inappropriate. This does not mean abandoning statistical analysis or quantitative rigor, however. Non-confirmatory studies can use other statistical tools, including exploratory data analysis (EDA) [31] and Bayesian statistics [32]. Unplanned sample augmentation is specifically problematic for *p* values; other statistical measures do not have the same problem (e.g., compare Figure 1 to Supplementary Feigure 2)[33, 34]. Therefore, in transparently non-confirmatory research, unplanned sample augmentation isn’t even N-hacking. If a sampling decision heuristic of the sort simulated here were employed, researchers need not worry too much about this producing an avalanche of false findings in the literature.

A problem in the Biology literature is that many non-confirmatory studies report performative *p* values and make “statistical significance” claims, not realizing that this implies and requires prospective study design. It is always improper to present a study as being prospectively designed when it was not. To improve transparency, authors should label non-confirmatory research as such – with no stigma implied. Journals and referees should not demand reporting of *p* values or “statistical significance” in such studies, and authors should refuse to provide them. Where to draw the boundary between approximately confirmatory and non-confirmatory remains blurry, however. My own opinion is that it is better to err on the side of classifying research non-confirmatory, and reserve null hypothesis significance tests and p values for cases where there is a specific reason a confirmatory test is required. **Broader Implications**

In the pursuit of improving the reliability of science, we should *question* “questionable” research practices, rather than merely denounce them [35-44]. We should also distinguish practices that are inevitably, severely misleading [45-47] from ones that are only a problem under specific conditions, or that have only minor ill effects. A quantitative, contextual exploration of the consequences of a research practice is more instructive for researchers than giving them a blanket injunction. Such thoughtful engagement can lead to more useful suggestions for improved practice of science, or may reveal that the goals and constraints of the research are other than what was assumed.

## Data Availability Statement

The code used to perform all simulations, the saved results of the simulation runs, the code used to generate the figures, and additional simulations and figures, are available on CodeOcean at https://doi.org/10.24433/CO.6897218.v1.

## Acknowledgements

I am grateful to Hal Pashler, Casper Albers, Daniel Lakens and Steve Goodman for valuable discussions and helpful comments on earlier drafts of the manuscript.

## Supporting Information Captions

Supplementary Table S1. Relation of the window width parameter w to the lower and upper cutoff p values defining the eligibility window, for the case of α=0.05.

Supplementary Table S2. N-hacking compared with alternative fixed-N policies.

Supplementary Figure S1. Data from the simulations shown in Figure 1, re-analyzed using log likelihood ratios instead of p values.

Appendix 1: Definitions of terms and variables as used in this paper

Appendix 2: A conservative bound on Type I Error Rate?

