## Supporting Information for "Is N-Hacking Ever OK? A simulation-based inquiry"

### Supplementary Materials

| $\alpha = 0.05$ | | | |
|---|---|---|---|
| $w$ | $P_{min}$ | $P_{max}$ | $\alpha_{procedure} \leq$ |
| 0 | 0.05 | 0.050 | 0.050 |
| 0.2 | 0.05 | 0.060 | 0.055 |
| 0.5 | 0.05 | 0.075 | 0.063 |
| 0.8 | 0.05 | 0.088 | 0.069 |
| 1 | 0.05 | 0.100 | 0.075 |
| 2 | 0.05 | 0.150 | 0.100 |
| 18 | 0.05 | 0.950 | 0.500 |
| 19 | 0.05 | 1.00 | 0.520 |

**Supplementary Table S1.** Relation of the window width parameter *w* to the lower and upper cutoff *p* values defining the eligibility window, for the case of $\alpha = 0.05$. If an interim *p* value falls between these cutoffs, the result is considered “inconclusive” or “promising”, and sample size is incremented, subject to some finite cap on the sample size. If an interim *p* value falls below the lower cutoff, sampling terminates with a decision of “significant”. If an interim *p* value falls above the upper cutoff, sampling terminates with a decision of “nonsignificant” (fail to reject the null) or “futile”. The case of $w = 0$ is equivalent to a fixed-N sampling procedure. Values of $w \leq 1$ are posited to be representative of informal heuristic decisions used by some researchers. Egregious N-hacking ($w > 2$, red) is incompatible with reporting p values.

| Procedure | <N> | $FP_0$ | Power | PPV |
|---|---|---|---|---|
| A. Fixed-N | 8 | 0.05 | 0.46 | 0.51 |
| B. Augmented | 9 | 0.12 | 0.78 | 0.42 |
| C. Fixed-N | 9 | 0.12 | 0.68 | 0.39 |
| D. Fixed-N | 9 | 0.19 | 0.78 | 0.31 |

**Supplementary Table S2** N-hacking compared with alternative fixed-N policies. **A.** Performance of a fixed-N procedure with $N = 8$, $\alpha = 0.05$. **B.** Performance characteristics of a constrained sample augmentation condition that had higher power but lower PPV: $N_{init} = 8, \alpha = 0.05, w = 5$. **C.** Fixed-N procedure where N and $\alpha$ are chosen to match the final sample size <N> (blue) and false positive rate $FP_0$ (yellow) of the procedure in B, showing that this is worse than B (lower power and PPV). **D.** Fixed-N procedure where N and $\alpha$ are chosen to match <N> and power (green) of procedure B, showing that this is worse than B (higher $FP_0$ and lower PPV).

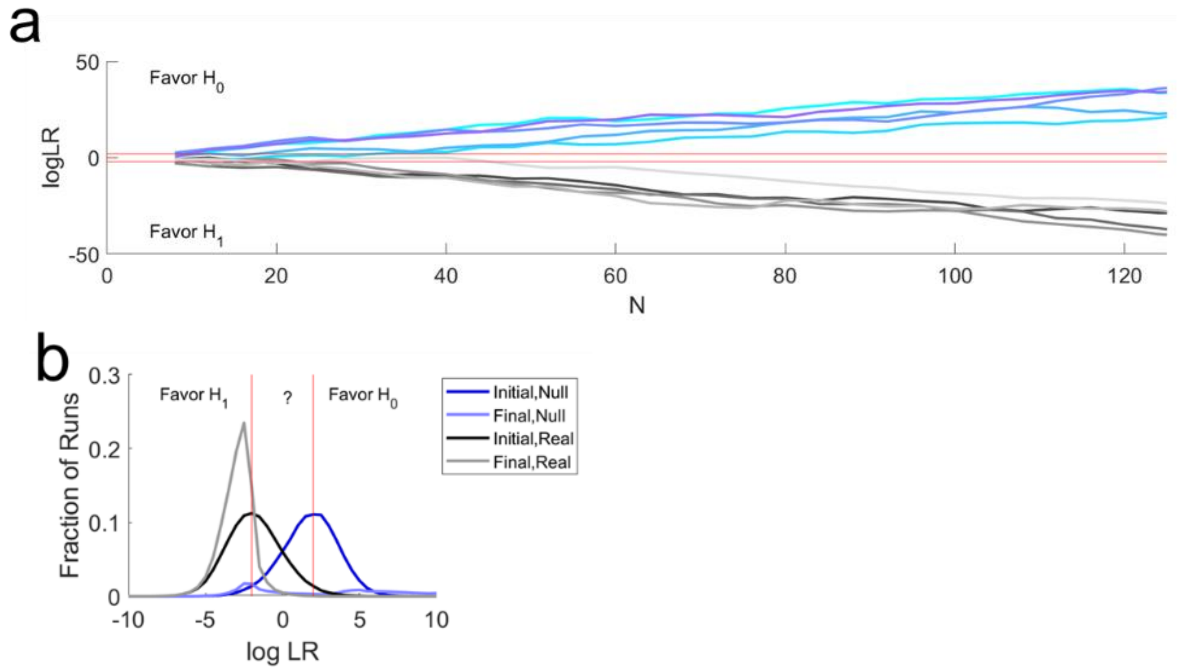


**Supplementary Figure S1. Data from the simulations shown in Figure 1, re-analyzed using log likelihood ratios instead of *p* values.** Here the null hypothesis ($H_0$, no effect) is compared to a specific alternative hypothesis ($H_1$, a positive effect of 1SD). The weight of evidence is measured by the $\log_{10}$ of the Likelihood Ratio (log LR) of the two hypotheses. A negative number indicates more evidence for the alternative (a real effect) and a positive number indicates more evidence for the null (no effect). A possible decision criterion is indicated by the red lines: accept the null hypothesis if $\log LR > 2$ (i.e., likelihood ratio 100:1 in favor of null), accept the alternative if $\log LR < -2$ (100:1 in favor of alternative), and consider the study inconclusive if log LR is between those thresholds. **a.** Evolution of log LR with sample growth, for five example runs from the egregious N-hacking simulation analyzed in Figure 1, in which the null hypothesis was true (shades of blue). Five example runs from experiments with a real effect of 1SD (shades of gray) are also shown. As the sample size increases, log LR fluctuates. But unlike the *p* value, which fluctuates randomly with no net trend, log LR *trends systematically* toward the null conclusion for null experiments, and systematically toward the real-effect conclusion for experiments with real effects. This matches experimentalists' intuition that having more data is always more informative. **b.** Distributions of final log LR values after egregious N-hacking (c.f. Figure 1b, distribution of *p* values of the same runs). For the null experiments, the vast majority of inconclusive cases (dark blue curve, area between the red lines) resolve to *negative* results (most of the area under the pale blue curve is shifted off the scale to the right). For the experiments with real effects, most of the inconclusive cases (black curve, area between the red lines) resolve to positive results (most of light gray curve is shifted to the left of the criterion for accepting $H_1$). It is controversial whether or not Bayes Factors are immune to N-hacking [S1-S4], however.

#### Appendix 1: Definitions of terms and variables as used in this paper

| | |
|---|---|
| Sample | A group of representatives randomly selected from a larger population and intended to represent it |
| Observation, data point | One of the individuals or representatives in a sample |
| $H_0$ Null hypothesis (no effect) | For example, in an independent sample *t*-test comparing samples from populations A and B, the null hypothesis is that the means of the groups are the same: $H_0: \mu_A = \mu_B$ |
| $H_1$ Alternative hypothesis (effect) | For the *t*-test example, the alternative is that means of the populations are not the same: $H_1: \mu_A \neq \mu_B$ |
| $N$ Sample size | In a fixed-N procedure: the number of observations in each group<br>In an incrementing procedure:<br>$N_{init}$ Initial sample size<br>$N_{incr}$ Number of observations added each time<br>$N_{max}$ Maximum sample size before stopping |
| $p$ Value returned by statistical null hypothesis test | The fraction of such experiments in which one would observe a difference at least as great as the observed difference, if in fact $H_0$ were true. |
| $\alpha$ Significance criterion | A criterion to reject $H_0$ only if $p < \alpha$ |
| $w$ Eligibility window | In the N-increasing procedure, defines how close to $\alpha$ a $p$ value must be to collect more data as follows:<br>$\alpha \leq p < (1 + w)\,\alpha$ |
| $FP_0$ False Positive Rate on the Null (Type I Error Rate) | For any procedure, probability of rejecting the null if the null is true: $FP_0 \equiv P(reject\ H_0 \mid H_0)$<br>For the fixed-N case in this paper, $FP_0 \equiv \alpha$<br>In simulations: the observed frequency of positive results when both samples drawn from the same distribution. |
| $E$ Effect size | The true difference in the means of the two populations being compared, expressed in standard deviations:<br>$E \equiv \frac{\lvert \mu_A - \mu_B \rvert}{\sigma}$ |
| $P(H_1)$ Prior probability of an effect | The probability $H_0$ is false, before considering the data. In simulations: fraction of experiments in which the samples were drawn from distributions with different means. |
| Power | The probability that a real difference will be found to be significant: $Power \equiv P(reject\ H_0 \mid H_1)$<br>Depends on $\alpha$, $N$ and effect size $E$ |
| $PPV$ Positive Predictive Value | The probability that an effect that was deemed significant is in fact real: $PPV \equiv \mathrm{P}(H_1 \mid reject\ H_0)$<br>Depends on $\alpha$, Power, and prior $P(H_1)$<br>Related to "False Positive Risk"[48, 49]: $FPR = 1 - PPV$ |

#### Appendix 2: A conservative bound on Type I Error Rate?

In some cases a strictly confirmatory study is needed to inform a high-stakes binary decision, in which case a prespecified sampling procedure is essential. In such cases researchers may still feel that pre-registering a fixed sample size is overly constraining, as they want the flexibility to abandon data collection early if results are not promising, or to continue data collection if results are very promising. Although there are many standard adaptive sampling procedures available, many researchers find them prohibitively complicated or confusing. Therefore it might be of some value to point out that **the procedure described, if pre-registered, would be entirely valid in a confirmatory setting**.

Although no formal proof has been provided, the simulated data strongly suggest that if one committed to the simulated decision rule formally in advance – without any multiple comparison correction for re-tests, as simulated – the following inequalities apply (dotted lines, Figure 4e):

$FP_0 < \alpha(1 + \frac{w}{2})$ for $N_{incr} \leq N_{init}$

$FP_0 < \alpha(1 + \frac{w}{4})$ for $N_{incr} = N_{init}$

These appear to be loose bounds; in many conditions the false positive rate falls well below this value. But they have the virtue of being trivial to calculate. For example: an N-increasing procedure with $w = 0.4$, $N_{init} = 10$, $N_{incr} = 10$, $N_{max} = 50$, would have a bound of $FP_0 < 0.0550$ by rule of thumb, compared to the simulation result of $FP_0 = 0.0541 \pm 0.0001$ (mean ± SD). The code provided [13] can be used to numerically estimate the false positive rate for any parameter combination. Using worst-case parameters $\alpha = 0.10$ $N_{incr} = 1$ $w = 9$ for $N_{init} = 128$ with $N_{max} = 256$ still did not exceed this empirical bound, despite allowing for up to 128 "peeks" at the data with no peeking penalty.

Thus, if one committed to using the procedure simulated in Fig 2A ($\alpha = 0.05$, $w = 1$) one could conservatively report $\alpha_{procedure} < 0.075$ regardless of $N_{init}$ and $N_{incr}$. Or one could choose a nominal $\alpha = 0.0\overline{33}, w = 1$ ($p_{max} = 0.0\overline{66}$) in the procedure to guarantee $\alpha_{procedure} < 0.05$. Or if a lab followed a general policy of collecting more data only if $p < 2\alpha$ ($w = 1$), they could conservatively correct for the possibility of unplanned sample augmentation by reporting an adjusted false positive rate of $1.5\alpha$ (for all their experiments, whether or not augmentation occurred in that case).

I have not done a quantitative comparison of the performance characteristics of this procedure to well-established adaptive sampling procedures, and I am making no claim that it would be especially efficient or optimal. I have not found this exact sampling procedure described in the statistical literature, although new adaptive sampling procedures are continually being proposed, and the one explored here belongs in the generally family of promising zone methods.
